# Feature selection strategies for drug sensitivity prediction

**DOI:** 10.1101/856013

**Authors:** Krzysztof Koras, Dilafruz Juraeva, Julian Kreis, Johanna Mazur, Eike Staub, Ewa Szczurek

## Abstract

Drug sensitivity prediction constitutes one of the main challenges in personalized medicine. The major difficulty of this problem stems from the fact that the sensitivity of cancer cells to treatment depends on an unknown subset of a large number of biological features. Although feature selection is the key to interpretable results and identification of potential biomarkers, a comprehensive assessment of feature selection methods for drug sensitivity prediction has so far not been performed. We propose feature selection approaches driven by prior knowledge of drug targets, target pathways, and gene expression signatures. We asses these methodologies on Genomics of Drug Sensitivity in Cancer (GDSC) dataset, a panel of around 1000 cell lines screened against multiple anticancer compounds. We compare our results with a baseline model utilizing genome-wide gene expression features and common data-driven feature selection techniques. Together, 2484 unique models were evaluated, providing a comprehensive study of feature selection strategies for the drug response prediction problem. For 23 drugs, the models achieve better predictive performance when the features are selected according to prior knowledge of drug targets and pathways. The best correlation of observed and predicted response using the test set is achieved for Linifanib (r=0.75). Extending the drug-dependent features with gene expression signatures yields models that are most predictive of drug response for 60 drugs, with the best performing example of Dabrafenib. Examples of how pre-selection of features benefits the model interpretability are given for Dabrafenib, Linifanib and Quizartinib. Based on GDSC drug data, we find that feature selection driven by prior knowledge tends to yield better results for drugs targeting specific genes and pathways, while models with the genome-wide features perform better for drugs affecting general mechanisms such as metabolism and DNA replication. For a significant group of the compounds, even a very small number of features based on simple drug properties is often highly predictive of drug sensitivity, can explain the mechanism of drug action and be used as guidelines for their prescription. In general, choosing appropriate feature selection strategies has the potential to develop interpretable models that are indicative for therapy design.

## Background

The ability to predict a response of a specific cancer type to a therapy is one of the main goals in precision medicine. Considering molecular features of cancer cells is crucial for mitigating heterogeneity and for tailoring the therapy to specific patients (1). The emergence of large scale high-throughput screening studies (2–6) have allowed researchers to develop computational models for drug response prediction from molecular profiles of human cancer cell lines or drug properties (7, 8). Although the inconsistencies and limitations of cell line data have been raised and extensively studied (9–12), these resources remain a vital tool for development of such models.

Arguably, the desired quality of computational models of drug response is not only their predictive performance, but also interpretability. To evaluate candidate drug efficacy on a specific patient’s tumor, many approaches apply black-box algorithms with a set of highly dimensional features as input. In clinical practice, the capability of extracting such hig-hvolume data from patients material is limited. Thus, there is a growing need of proper identification of concise, limited subset of features, or biomarkers, that are most informative of drug response. Therefore, strong emphasis should be put on feature selection approaches for drug sensitivity prediction. Despite its paramount importance, no systematic assessment of feature selection strategies in the task of drug response prediction was so far performed.

The problem of drug response prediction has been approached by a wide spectrum of linear and non-linear machine learning algorithms, including regularized linear regression, k-nearest neighbors (KNN), support vector machines and random forests (13–18). Multitask learning was proposed to improve drug sensitivity prediction by pooling information learned for different drugs (15, 19). Finally, a number of highly predictive kernel-based multi-view and multi-task models were introduced for drug sensitivity (20–22). Although highly predictive, the multi-task and multi-view approaches suffer from low interpretability. As a remedy, a multi-task learning approach based on a Bayesian model for collaborative filtering was proposed (23), which allows for identifying general interactions between features of the drugs with features of the cell lines. For example, it gives insights in the form of ”activation of pathway Y will confer sensitivity to any drug targeting protein X". This approach, however, does not directly address the crucial need of identifying biomarkers for specific drugs.

For that aim, data-driven, automatic techniques of feature selection were applied (17, 22, 24). Generally, the problem of identifying the optimal subset of features is intractable (25). Data-driven feature selection thus proceeds either as a heuristic search over the space of feature combinations, or is embedded directly in the learning algorithm by imposing sparsity of parameters associated with the features via regularization. Although these methods can achieve good predictive performance and deal with the curse of high data dimensionality (17), feature importance estimates and selection might not always be accurate and stable, especially in vastly high-dimensional data and in the presence of correlation between features (26). Stability selection was proposed to mitigate this problem when regularized regression is applied (27), but it still comes without the guarantee to choose the most biologically relevant predictive features.

Drug prediction approaches largely differ with respect to the type of features that they model. Among the molecular data feature types which characterize the cancer cell lines, gene expression was assessed as the most informative, with remaining types such as mutation or copy number data bringing limited predictive power (13, 14). Accordingly, genome-wide gene expression is the most common choice in the case of models utilizing single data type (7, 17, 22, 23, 28). Other studies reported that in some cases gene expression alone might not be sufficient, especially in a cancer- or drug-specific setting (29, 30). Importantly, expanding the feature space related only to cancer cell lines’ biology with drug-related properties was shown to improve predictive performance (15, 21–23, 30). The predictive drug-specific features may be related to their chemical properties, such as compound structure (15, 30), their known primary targets or pat-hway activation (21, 22).

Here, we utilize the knowledge regarding drug targets and their mode of action to select plausible features describing the cancer cell lines. This drug-related prior knowledge is thus used to directly limit the initial feature space, rather than first expanding it and next using data-driven selection techniques to narrow it down. We argue that this approach for feature selection in combination with common regression techniques can provide a simple and highly interpretable model without losing the predictive performance characteristic for models starting from high-dimensional data. In fact, the direct utilization of prior knowledge is the number one strategy recommended for feature selection according to the classics in machine learning (25). It was however, never exploited in the task of drug response prediction. We assess this methodology in a systematic fashion for a broad spectrum of anti-cancer compounds, integrating multiple data types and comparing the results to the baseline models utilizing genome-wide gene expression data and data-driven feature selection techniques. On top of that, we evaluate gene expression signatures as the means of dimensionality reduction of the transcriptomics data and evaluate their predictive power in this context. This comprehensive analysis pin-points a set of drugs for which easily interpretable, informative, small sets of features can be identified.

## Methods

### Analyzed dataset

The analyzed dataset was acquired from the Genomics of Drug Sensitivity in Cancer (GDSC) (3) database. A total of 251 compounds were included in the analysis. Each was assigned one of 24 classes of target path-ways, defined by the GDSC.

The total set of samples consisted of 983 cancer cell lines originated from 13 tissue sites. The available data types for describing the cell lines included: gene expression (17737 features), coding variants (310 features), copy number variants (CNV, 425 features) and tissue type (13 features). Coding variants and copy number variants were represented as binary calls determining the presence or absence of a variant in a given gene or segment, respectively. We have dummy encoded the tissue types resulting in 13 distinct binary features for every cell line. All biological input data were acquired directly from the GDSC resource.

GDSC provides two types of metrics representing the drug efficacy: half maximal inhibitory concentration (IC_50_) and area under the dose-response curve (AUC). Since in our analysis we did not observe significant differences in predictive performance when using one metric in favor of the other, we picked AUC as our single target variable.

### Predictive algorithms

We employed two common machine learning algorithms in order to predict the AUC values: elastic net linear regression and random forest regression. We implemented both methods using Python3 scikit-learn 0.19.2 library (31). See Supplementary Methods for descriptions of the algorithms and implementation details.

### Feature selection

With a total of 18485 biological features that can be used to describe the cancer cell lines, the analyzed dataset is very high-dimensional. In contrast, the number of samples is in the order of hundreds, which poses the danger of overfitting. This might especially be the case when considering all available genome-wide information regardless of the drug being modeled. Here, we investigate different feature selection methods to mitigate this problem. These approaches can be divided into two groups: biologically driven and automatic, data-driven selection methods.

#### Biologically driven feature selection

##### Features based only on drug targets and tissue type, shortly only targets (OT)

In the most restricted feature space, we included only predictors corresponding to the direct targets of the drugs, as well as tissue type. Drug targets information was derived directly from GDSC. As an additional resource, we used DrugBank (32) database, assigning targets for 88 matched compounds. For each drug target, we included features representing the target gene’s expression, coding variant and copy number variation. In the case of copy number data, a given genetic feature was incorporated if the corresponding segment included at least one of the drug target genes. We only considered drugs with explicit gene targets annotation in GDSC or DrugBank and for which at least one feature in addition to the tissue type was available in the data. These conditions were met for 184 compounds. Applying two regression algorithms for each drug resulted in 368 separate models.

##### Set of features based on drug targets, tissue type, and target pathways, shortly pathway genes (PG)

In this approach, we included features related to genes that belonged to the same signaling pathway as the set of target genes. Pathways information was derived from Reactome (33, 34) database (version 66 accessed on October 2018). For each compound, first its target set was derived, followed by finding all pathways which included at least one of the given targets. The total set of considered genes was then computed as the union of all members of the found pathways. Lastly, corresponding gene expressions, coding variants, copy number variants and tissue types were extracted to create the final feature set. The drug targets and pathway information was available for 186 drugs, producing 372 models.

##### Sets of features resulting from addition of gene expression signatures, shortly OT + S or PG + S

Gene expression signatures can explain the activation level of complex biological phenomena in the investigated cell lines. Here, we refer to a gene signature as a set of genes related to a certain known biological phenomenon that can be deduced from cancer gene expression data (Supplementary Table S1). For each signature *S* with *i* genes, we calculated two scores. The first characterizes the coherent expression and the second estimates the activation level of *S*. Given a gene expression matrix for *S* in *n* samples (*X*^*i×n*^), the previously described coherence score (CS) (35), is calculated as the mean pairwise Pearson correlation between all columns of *X*. Therefore, a strong negative or positive correlation between all genes in *S* is indicated by CS values close to −1 and 1, respectively. The activity of *S* (i.e. the signature score) for each sample is calculated by first *z*-scoring the gene expression values across samples, followed by averaging the resulting *z*-scores across genes. Here, we calculated the signature scores using the cancer cell line expression data provided by GDSC. We set the threshold for a significantly coherent activation of *S* to CS(*S*) ≥ 0.1, resulting in 128 signature features. The OT + S set contains features based on target genes, signature scores and tissue type. The PG + S set contains target genes, pathway genes, signature scores and tissue type. Applying two regression algorithms for each drug resulted in 740 separate models.

##### Set of features based on genome-wide gene expression, shortly genome-wide (GW)

Finally, we constructed a feature set based exclusively on the expression of 17737 genes as features. We evaluated this feature set for 251 drugs in total, resulting in 502 different models.

#### Data-driven feature selection, shortly GW SEL

In addition to feature pre-selection based on drug properties and biological relevance, we also evaluated automated feature selection algorithms in application to genome-wide expression data. We used two techniques, based on linear and non-linear methods. First, stability selection, which uses lasso regression on multiple bootstrap samples in order to choose robust features (27). Such selected features were next passed as input for elastic net models (further referred to as *GW SEL EN*). For the second technique, feature importance estimates derived directly from random forest were used. These features were then used for random forest regression models (*GW SEL RF*). For more detailed description of both techniques, see Supplementary Methods.

##### Prediction framework

For every feature space setting, we performed regression separately for each drug. The modeling process per single compound is described in Fig. 1. We first extracted the data associated with a particular drug and corresponding screened cell lines. We then randomly split the data into a training and a test set, with 0.3 of the data included in the test set. For each compound, we employed both elastic net and random forest regression. In order to find the best model, we performed randomized search over hyper-parameters grid, testing 30 parameters combinations. Each combination was evaluated using 3-fold cross-validation on the training data. Afterwards, the model with the best hyper-parameters was trained on the whole training set and applied to the input data in the test set. In order to improve robustness, the whole process was repeated five times for different train-test data splits and test results were averaged.

**Fig. 1.**
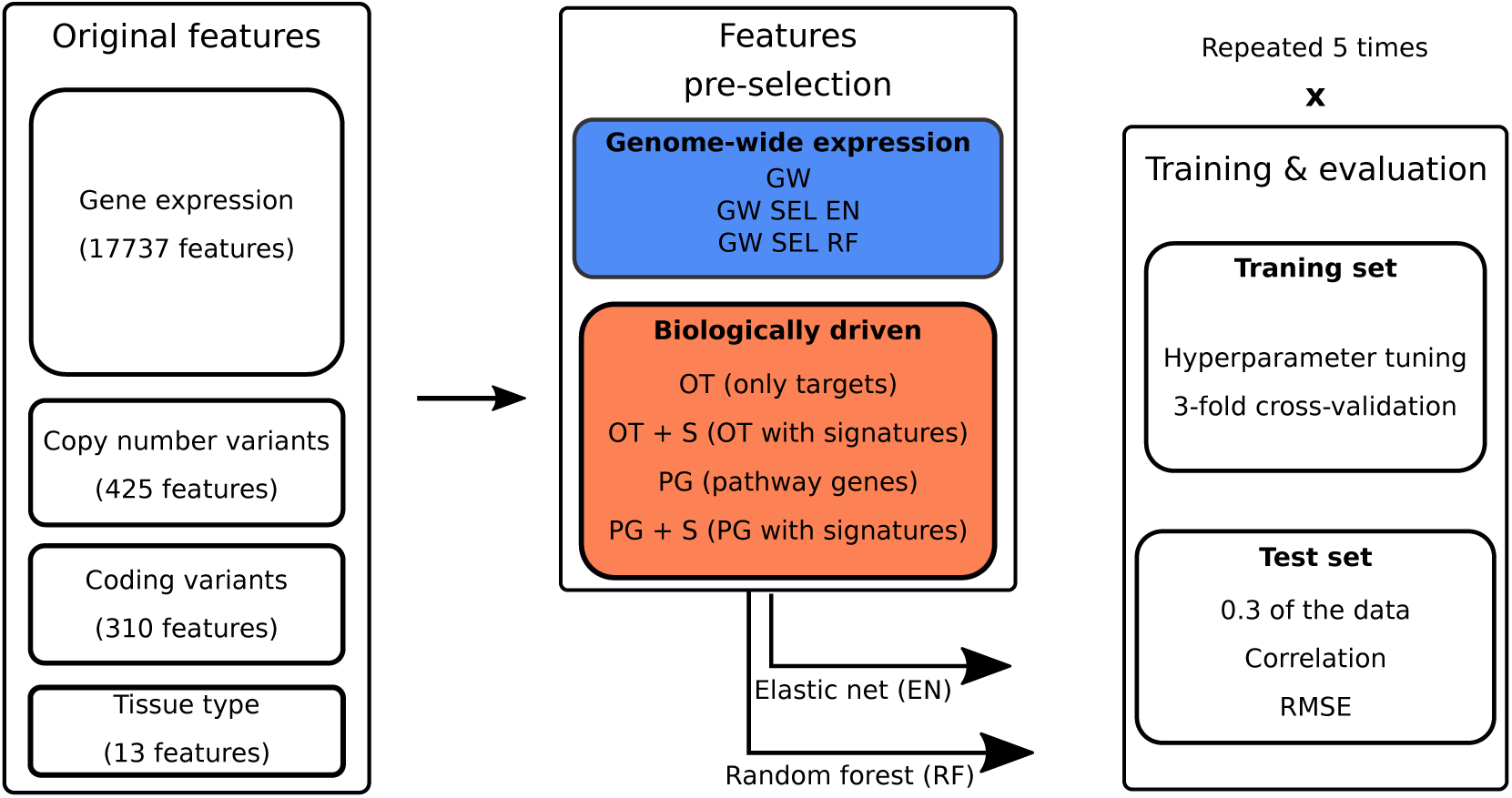
Flowchart describing the modeling framework for a single compound. Abbreviations: GW – genome-wide, PG – pathway genes, OT – only targets, EN – elastic net, RF – random forest, SEL – automated feature selection, S – gene expression signatures. For every feature space, we performed modeling separately for each drug. We randomly split the corresponding data into training and test set, with 0.3 of the data included in the test set. We used 3-fold cross-validation on the training data for hyperparameter tuning and evaluated the best model on the test set. The whole modeling process was repeated five times with different training/test set data splits.

##### Model evaluation

During cross-validation tuning, we used *Mean Squared Error* (MSE) as a scoring metric for best hyperparameters search. Although MSE is suitable for evaluation of different models within one compound, it is not reliable when comparing results across diverse drugs because of differences in corresponding AUC distributions. Furthermore, when a given target variable distribution has little variation, one can achieve a reasonably low MSE just by predicting the mean of a target variable. In order to avoid this problem and identify the models which performed well, we used *Relative Root Mean Squared Error* (RelRMSE), which is normalized in such a way that the score of 1 corresponds to a dummy model which always predicts the mean of target variable in the training data (see Supplementary Methods). The use of RelRMSE allowed us to distinguish drugs for which predictive algorithms could not outperform the dummy model, meaning that for those compounds no actual learning occurred.

In order to make further assessments and comparisons between compounds, we used Pearson correlation coefficient with the response AUC in the test set as a performance metric. As stated in a previous section, the recorded results for each method were averaged over five modeling procedures that were performed with different data splits.

## Results

### Models with genome-wide feature have larger feature sets and more samples than the models with biologically-driven features

The median numbers of input features were 3 and 387 for only targets and pathway genes feature sets, respectively (Fig. 2a). The input features were further expanded by including 128 gene expression signatures. In the case of methods based on automated feature selection, the optimal number of features, *k*, is shown. The median *k* are 70 and 1155 for random forests and for stability selection, respectively. All foregoing values constitute a drastic decrease in comparison to the number of 17737 genome-wide input features.

**Fig. 2.**
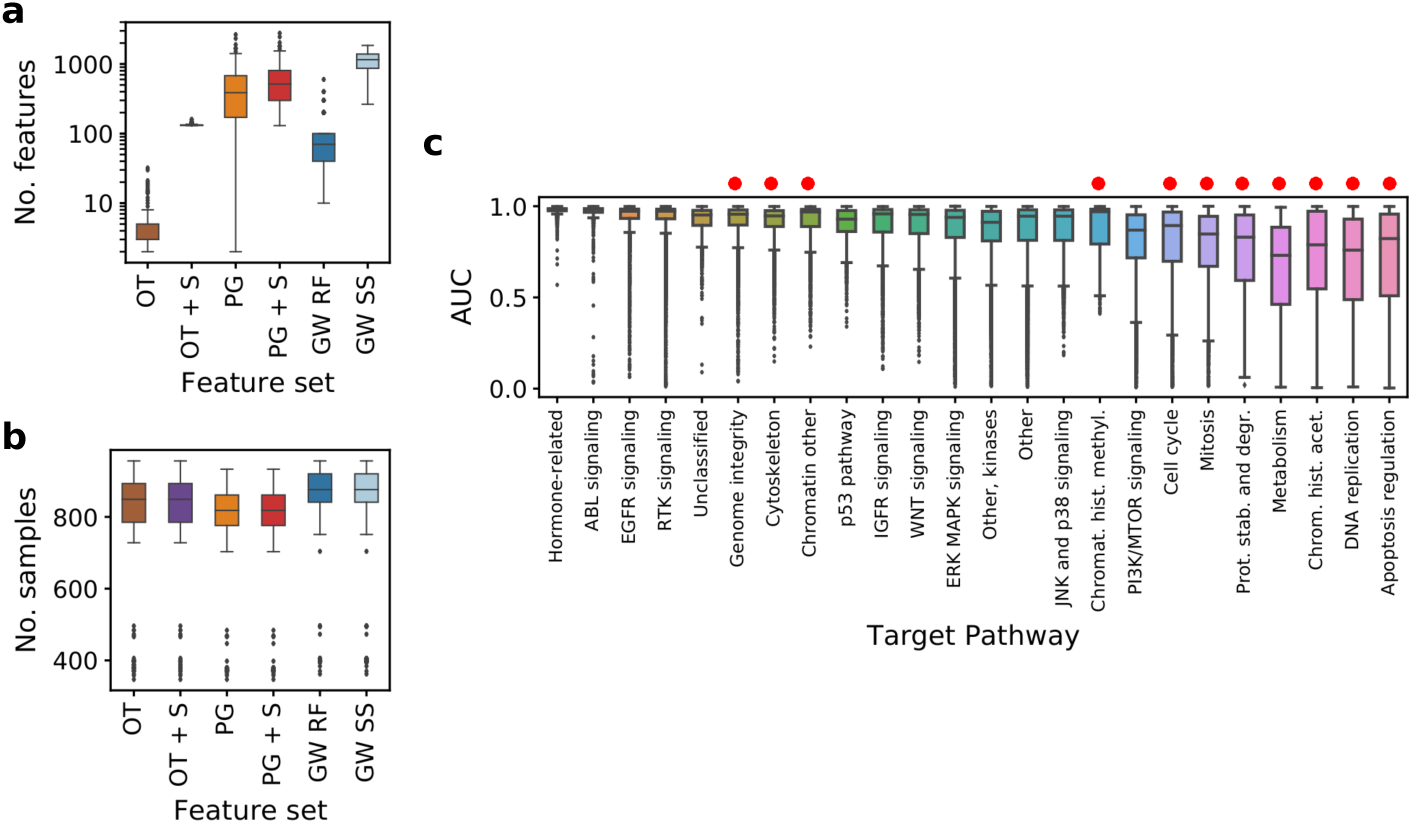
Models’ properties and overall results summary. (**a**) Number of input features across compounds in different methods. For genome-wide models, number of features was 17737 for each drug. Vertical axis uses log scale. (**b**) Number of samples across compounds in different methods. Abbreviation SS refers to stability selection (Methods). (**c**) AUC values grouped by target pathway of the drug, raw data from GDSC. Target pathways are sorted by interquartile range of the AUC values. Pathways corresponding to more general cell mechanisms are marked wit red dots. See Fig. 1 for abbreviations.

The number of samples for each drug also slightly differs for only targets and pathway genes feature sets, since for some cell lines the coding variants or CNV information are not available (Fig. 2b). This results in a lower number of samples for models with biologically driven features, with the median of 849 for only targets and 818 for pathway genes feature sets, compared to 876 for genome-wide expression features.

### AUC distributions are different across compounds, tend to have low variance for drugs targeting specific genes and pathways and high variance for drugs targeting general cellular mechanisms

The distribution of AUC values varies significantly among compounds with different target pathways (Fig. 2c). The median AUC value per target pathway ranges from 0.98 for hormone-related drugs to 0.73 for compounds targeting metabolism pathways. The smallest variation of AUC is observed for drugs targeting the hormone-related pathways. The largest AUC variation is observed for the apoptosis regulation pathway. The AUC for drugs targeting general mechanisms, such as DNA replication or metabolism, tends to have larger variance, which means their sensitivity is easier to model.

### The per-drug results show the importance of comparing to a dummy model and that different feature selection strategies are best suited for different drugs

Since RMSE measures the level of model error, and correlation measures the model agreement with the test set, both large (1-RMSE) and high correlation should coherently indicate a high model performance. The negative relation between (1 - RMSE) quantity and correlation, however, confirms the fact that raw RMSE is not a good metric for performance comparison between compounds (Fig. 3a; Methods). Instead, correlation achieved by the model increases with the modeled AUC variance (Fig. 3b).

**Fig. 3.**
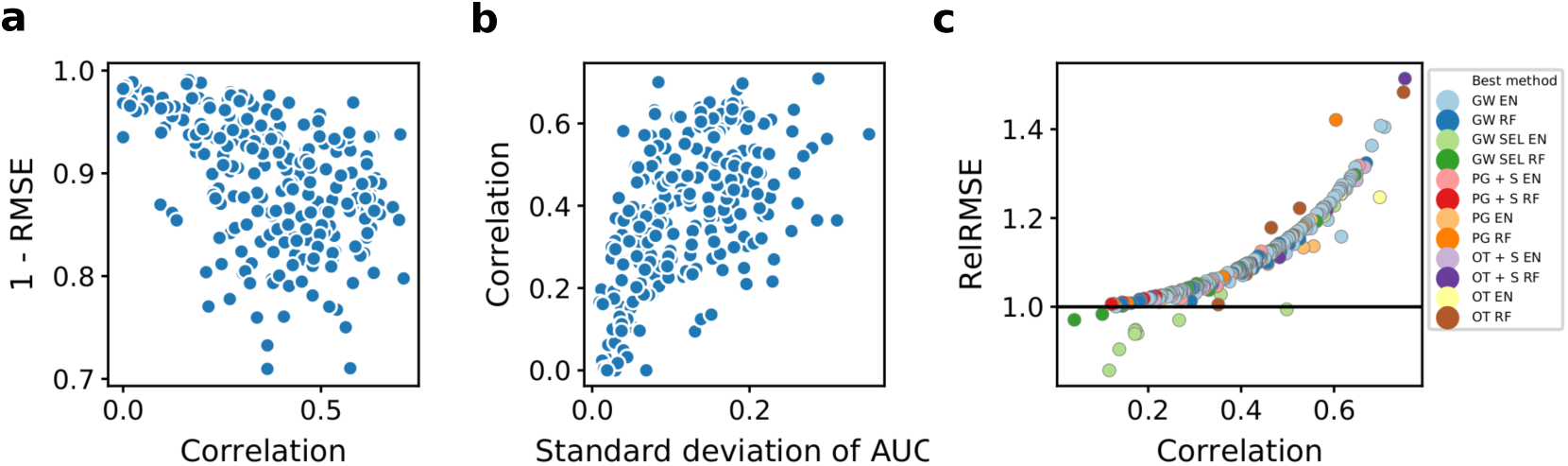
Predictive performance for all of the analyzed drugs. (**a**) 1 -RMSE vs correlation per drug, obtained by elastic net using genome-wide gene expression data as predictors. (**b**) Correlation vs. standard deviation of true AUC for all cell lines screened for a given drug, correlation obtained by genome-wide elastic net. (**c**) RelRMSE vs. correlation obtained by the best model for a given drug. Each point represents a single drug. For each of them, corresponding best performance was determined using correlation as a metric. Colors represent models with feature set which obtained the best performance for a given drug. Horizontal line at 1 represents the baseline RelRMSE score. See Fig. 1 for model abbreviations.

Both these facts support that RelRMSE is a better performance measure than raw RMSE (Fig. 3c; Methods). Indeed, RelRMSE grows with the correlation. Importantly, for some drugs, the best performing models fail to achieve the baseline RelRMSE score of 1 or are very close to 1 (Fig. 3c). Further inspection of these models reveals that they can capture only the mean AUC, since the modeled AUC distribution does not have enough variation. In total, there were 19 of such compounds and these were excluded from further analysis.

It is apparent from Fig. 3c, that for most of the drugs, the best suited method is modeling using genome-wide features and elastic net. However, this is not the case for compounds with the top corresponding modeling performances, as the two best correlation scores are achieved by models with biologically driven feature space. These two compounds are Dabrafenib and Linifanib, both with correlation of 0.75, for models with feature spaces: only targets genes with gene expression signatures and only targets genes, respectively. In terms of performance, they are followed by Trametinib (correlation 0.71) and Alectinib (correlation 0.70), both scores being achieved by genome-wide methods. In general, as we consider more top performances, the frequency of genome-wide methods among them increases, although they are not as highly represented when looking at the small group of absolute best scores.

### The difference in predictive performance of biologically driven versus genome-wide models is small, despite using significantly less input features

In general, genome-wide feature set combined with elastic net (GW EN) emerges as the best model with the median correlation of 0.39 (Fig. 4a). However, models with biologically driven feature spaces perform very similarly, (excluding only targets (OT) approaches), with the best median correlation of 0.37 produced by models employing target pathway genes features combined with gene expression signatures and elastic net (PG + S EN). Furthermore, the difference in median performance was negligible between genome-wide random forest (GW RF, with 17737 features) and genome-wide random forest with automated selection (GW SEL RF, with 70 features on average). This suggests that for many compounds, most gene expression features do not have significant power in predicting drug response. The spread in performance (defined as the difference between the maximum and the minimum value) reaches over 0.6 for all of the methods, suggesting that each drug should be approached individually in terms of modeling.

**Fig. 4.**
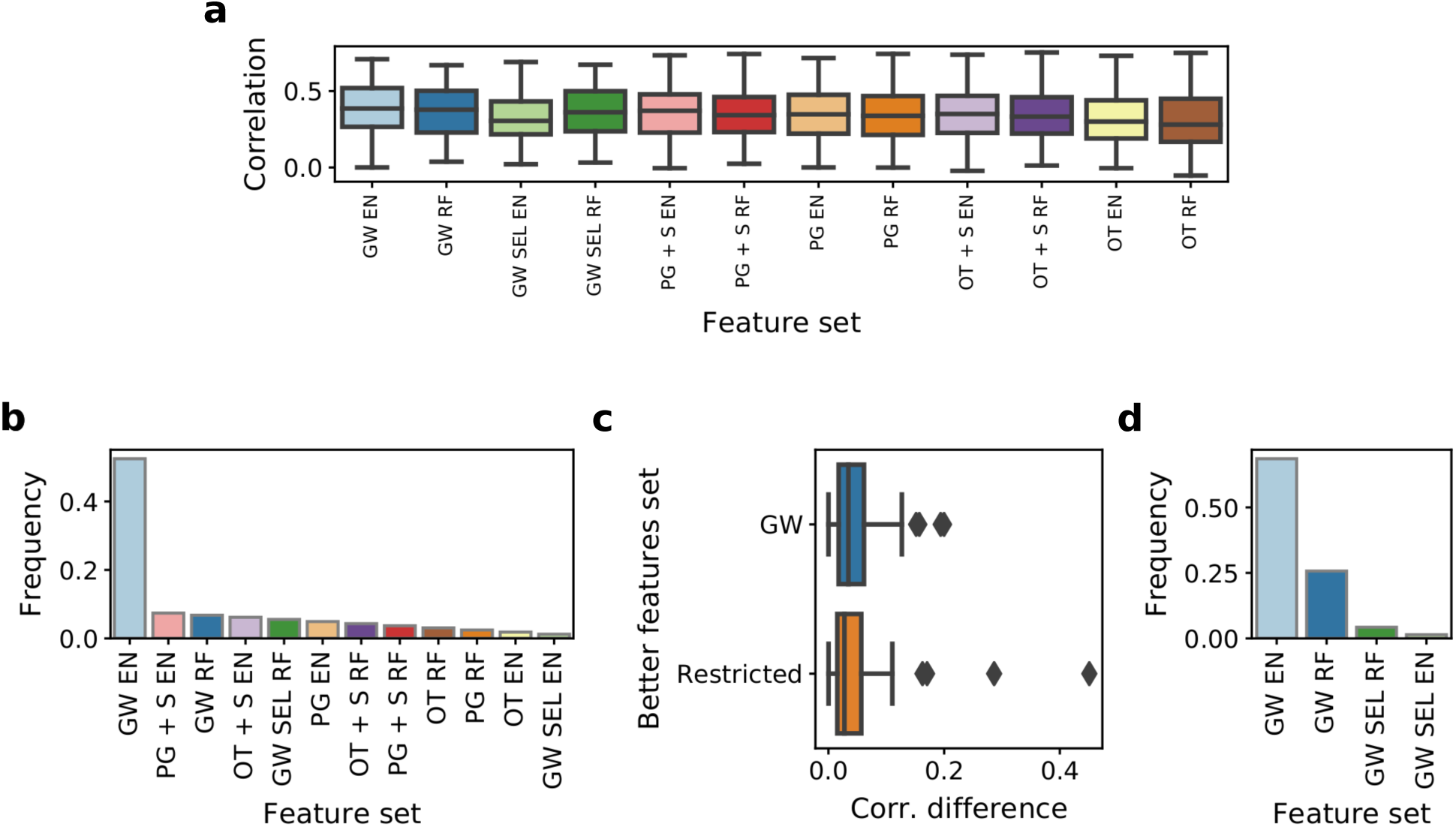
Frequencies of all applied methods among best models per drug. (**a**) Correlation of AUC predictions with the true AUC values in the test set across compounds in methods with different feature spaces. Results are shown for 175 drugs which were common across all applied models. (**b**) Model frequencies for compounds for which all methods were applied. (**c**) Differences in correlation between best model per drug overall and best model from the other class. Two cases are shown – genome-wide and biologically driven feature sets. (**d**) Model frequencies among best models for compounds where models with biologically driven could not have been applied. See Fig. 1 for abbreviations.

The standard, genome-wide model achieves the best performance for over half of considered drugs (Fig. 4b). However, for many of these cases the correlation difference between the best genome-wide model and the best model with biologically driven features is not significantly large, with the median of only 0.034 (Fig. 4c). The reverse is also true, with median correlation difference between the best biologically driven model and the worse genome-wide model 0.028. Despite a drastic reduction in feature space, the biologically driven models based either on only targets or pathways yield the best modeling performance for 23 drugs, outperforming all other models including the genome-wide approach. For further 60 drugs, the best models have feature space expanded with expression signatures. Noticeably, there are also 15 cases where data-driven feature selection helps to produce better performance with much smaller subset of the original feature set (Fig 4b, d).

### Predictive performance using different feature selection strategies depends on drugs’ target pathways

Next, we investigate the general tendencies concerning which feature selection is particularly better suited for modeling drugs targeting specific pathways. To this end, we compare the overall performance of biologically driven feature selection as one group to the baseline of genome-wide features and the genome-wide features with automatic selection as another (Fig. 5a), for different target pathways. Genome-wide models achieve better performance in 15 out of 24 pathways in total, however, the difference is statistically significant in only four of them (at 0.05 significance level): DNA replication, metabolism, apoptosis regulation pathways and a group of pathways referred to as "other". This indicates that these models capture a broad mechanism of action of the corresponding drugs. Conversely, the target pathways for which the models with biologically driven features most notably outperform models with genome-wide features include ABL, IGFR and EGFR signaling pathways, although these results are not statistically significant due to small sample sizes. The models with biologically-driven features perform also better for the hormone-related pathway, but overall the modeling performance is bad in this case and we do not consider this result reliable. In summary, compounds with specific signaling target pathways seem to benefit more from the initially restricted feature space.

**Fig. 5.**
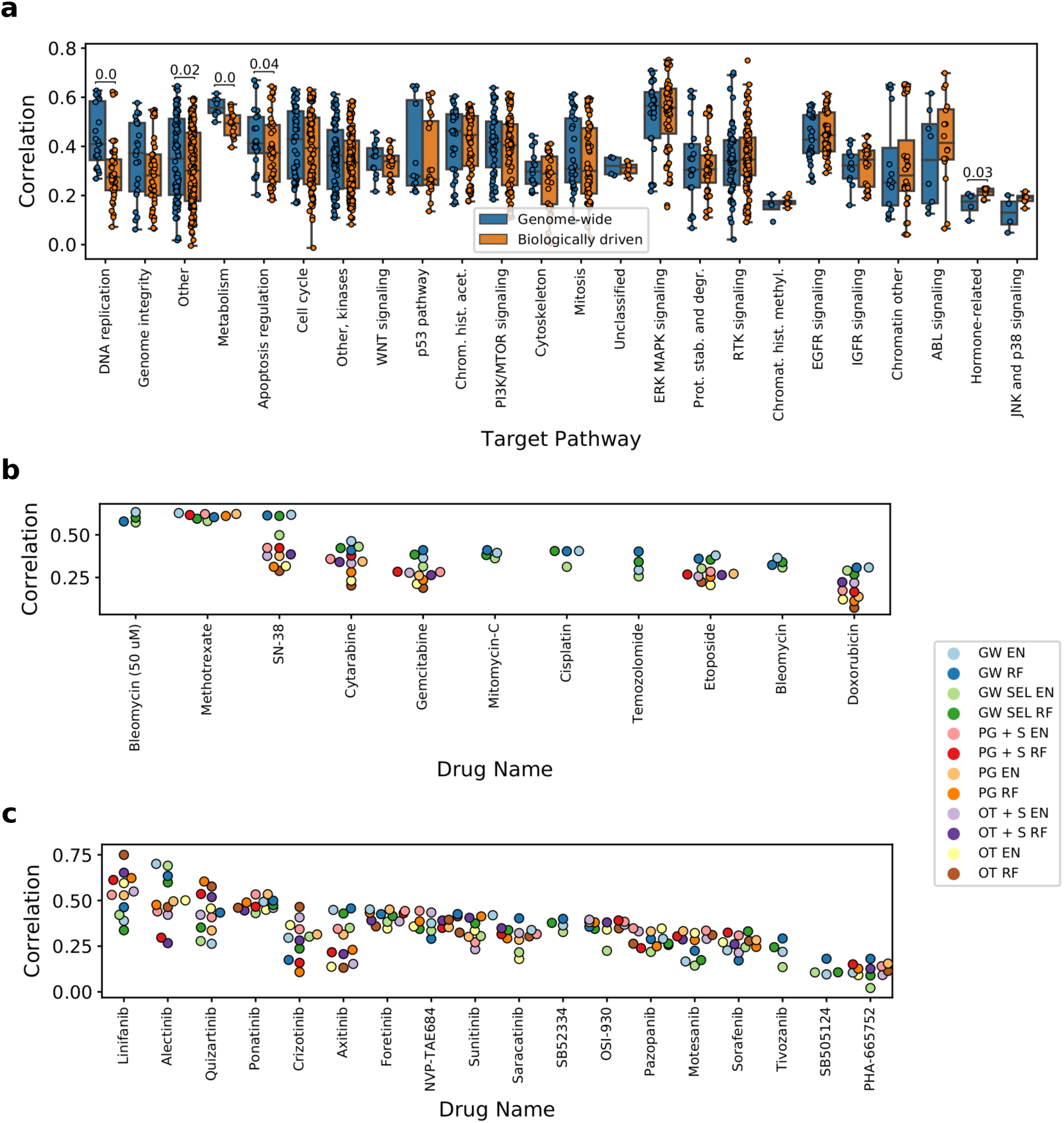
Predictive performance in relation to compounds’ target pathway. (**a**) Correlation with the test set grouped by pathways. Methods were classified into two groups – one that uses genome-wide feature space, and one with biologically driven feature space. Numbers displayed represent p-values of the significance test for difference. Lack of number means no statistical significance at 0.05 significance level. (**b**) Predictive performance for drugs with DNA replication target pathway. (**c**) Predictive performance for drugs with RTK signaling pathway. See Fig. 1 for model abbreviations.

We next inspect in detail the results for distinct drugs coming from DNA replication and RTK signaling pathways, respectively (Fig. 5b and c). Among the drugs targeting the DNA replication pathway, Bleomycin, Methotrexate and SN38 exhibit good modeling ability with the genome-wide features. However, in case of Methotrexate similar performance is achieved also by methods with biologically driven feature space, contrary to SN-38. Conversely to DNA replication pathway, among the drugs targeting the RTK signaling pathway the best result is more often produced by biologically driven features, with most noticeable cases of Linifanib and Quizartinib. In contrast, Alectinib exhibits good modeling performance exclusively with genome-wide approaches. In general, although the above described general tendencies apply, information about drug’s target pathway alone seems to be insufficient to clearly tell which feature space is the most suitable for predicting it’s response, with the potential exception of the DNA replication pathway.

### Gene expression and mutations constitute the most predictive feature types

In order to assess which feature types are most informative of drug response, we consider such models with biologically driven feature space, which use all five available data types (Fig. 6). To make results more robust, we consider only top 50 drugs in terms of corresponding modeling performance achieved by the biologically driven feature sets, resulting in worst considered model’s correlation of 0.47. Next, we extract top *k* most predictive features in each model and record the frequencies of particular data classes among them. Results confirm the fact that gene expression is the most predictive feature type, although mutation (coding variant) and tissue type are also important, especially for drugs designed to target specific cancer type with a particular mutation. In contrast, copy number variants seem not to incorporate much useful information. The relative effect of gene expression data increases with number of considered most predictive features, but this is expected given that this category is the most frequent of all available data types overall. Finally, the high frequency of gene expression signatures among the top predictive features implies that the signatures can act as good representatives of genome-wide information.

**Fig. 6.**
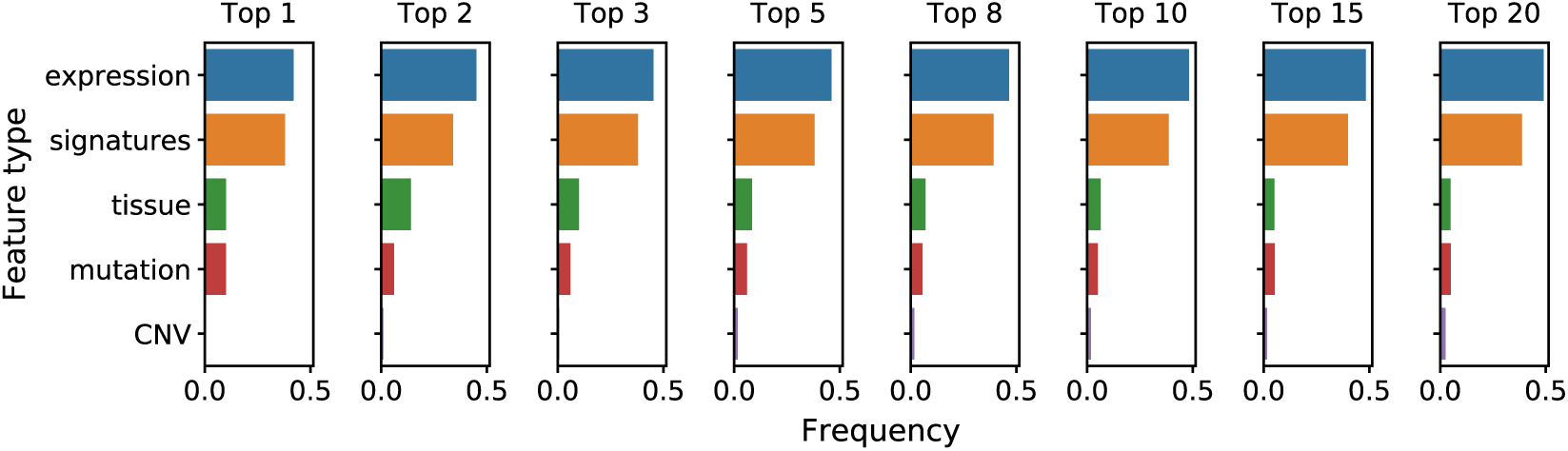
Frequencies of considered feature types among top *k* most predictive features. Feature importance coefficients were extracted from top 50 drugs in terms of modeling performance using methods with biologically driven feature space.

### Feature selection enables interpretation of the mode of action and pin-pointing biomarkers for the best modeled drugs

We further focus the analysis on ten drugs of most interest (Fig. 7), based on two simple criteria: top modeling performance achieved by all of the feature selection methods, or distinctly better performance achieved by one of the methods’ class (genome-wide or biologically driven) in comparison to another. In five of those compounds the best result is produced by models with the genome-wide features, whereas another five are better modeled with biologically driven features.

**Fig. 7.**
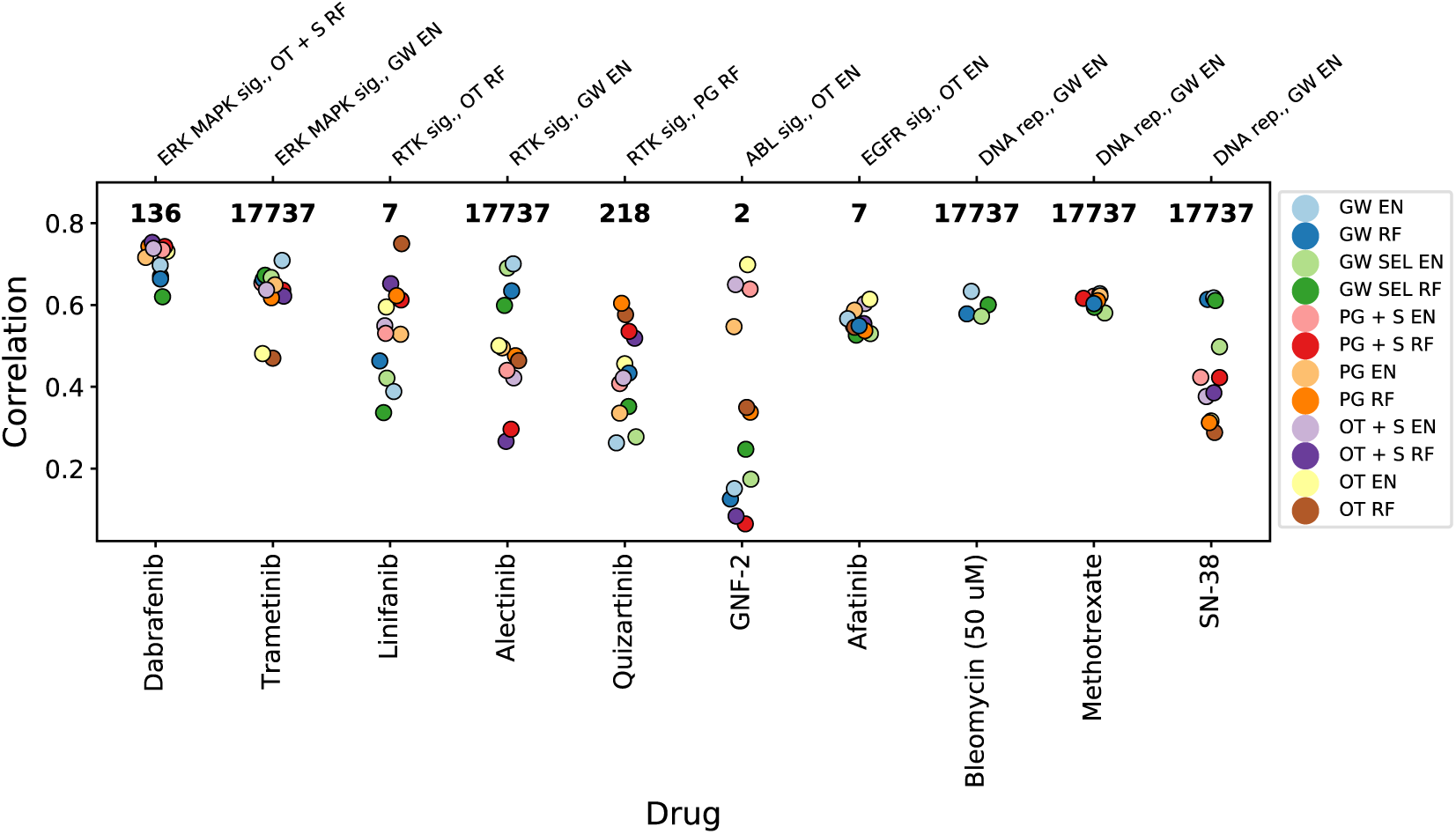
Results for specific compounds exhibiting good ability to model with one or all of the methods. Displayed numbers represent number of features which was used by best performing model for a particular drug. Top horizontal axis show compounds’ target pathways along with model which achieved the best modeling result. See Fig. 1 for model abbreviations.

From all analyzed drugs, Dabrafenib emerges as the compound which is the easiest to model. The highest correlation of 0.75 is achieved by the model with OT + S RF features in the biologically driven class, and performance of other approaches is only slightly worse (Fig. 7). This overall good modeling ability with the OT + S RF features could be explained by two factors. First, the AUC distribution corresponding to Dabrafenib is well-diversified, with relatively many cell lines sensitive to treatment (Fig. 8a), which leads to better modeling performance (compare Fig. 3b). Second, the relative effects of the selected features are in excellent concordance with the Dabrafenib’s pharmaceutical properties. The most predictive feature – mutation in BRAF oncogene (Fig. 8a) – and the second most predictve feature – the BRAF gene expression signature – well agree with the design of Dabrafenib as the BRAF inhibitor. Interestingly, the feature corresponding to BRAF gene expression alone ranks lower, 28 among 136 features for the best OT + S RF model and as low as 15817 among 17737 features for the GW EN model in terms of predictive power. Finally, in concordance with Dabrafenib’s intended use in treatment of BRAF mutation-positive melanomas and lung cancers (36, 37), the skin tissue feature is the third most predictive one for the best OT + S RF model.

**Fig. 8.**
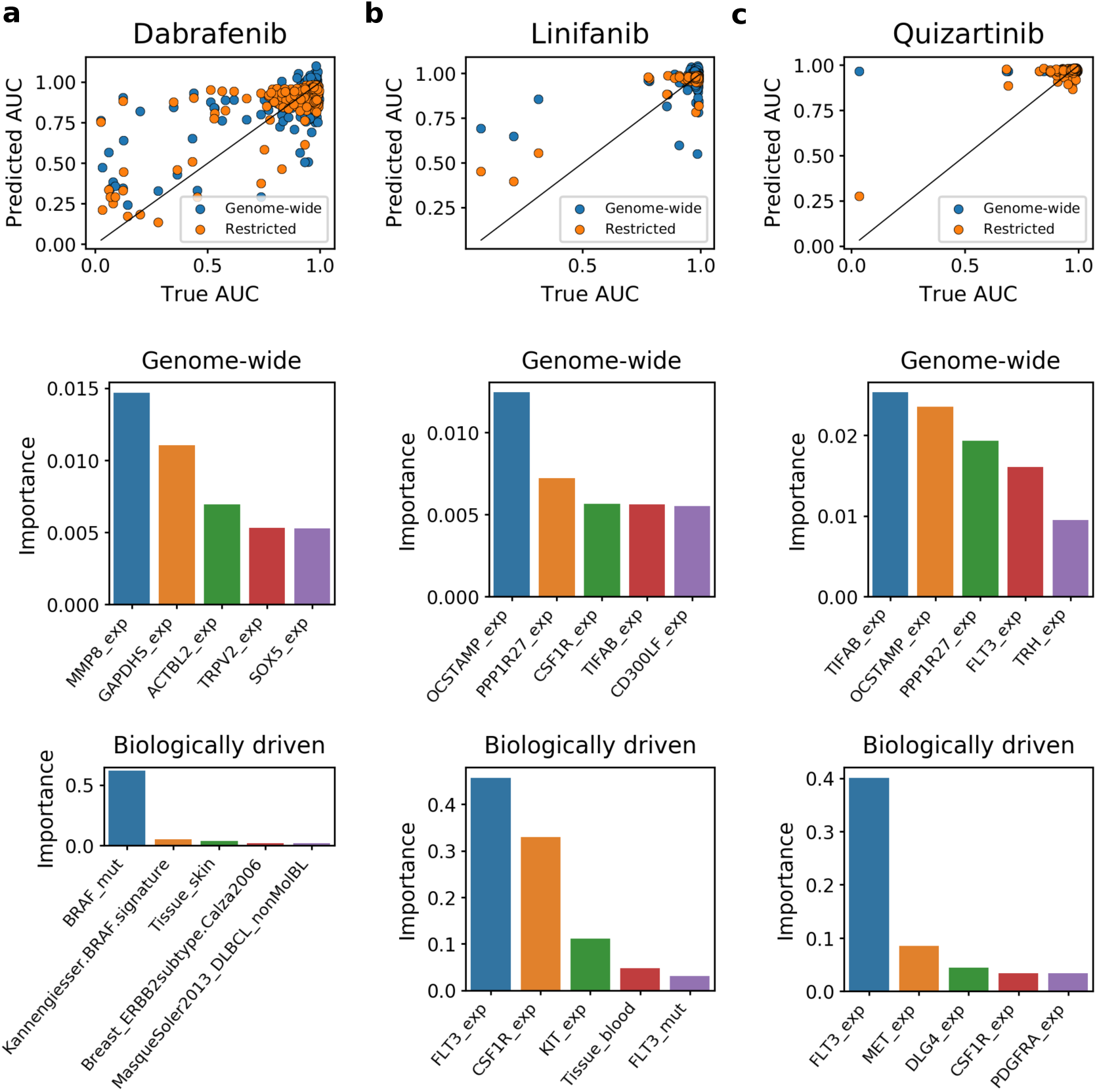
Predicted vs actual AUC values and most predictive features for (a) Dabrafenib, (b) Linifanib and (c) Quizartinib. Top panels show predicted vs actual AUC values when both biologically driven and genome-wide models were trained and tested on the same sets of samples. The biologically driven models correspond to best suited feature set for each drug: OT + S RF for Dabrafenib, OT RF for Linifanib and PG RF for Quizartinib. Middle and bottom panels present top 5 most informative features when fitting the model with genome-wide data (middle) and biologically driven feature space (bottom).

In the case of Linifanib, the best result (0.75 correlation) is accomplished by using only 7 features related to the drug’s targets (OT RF model, Fig. 7), which significantly outperforms the genome-wide models. Linifanib is an inhibitor of FMS-like tyrosine kinase 3 (FLT3) and vascular endothelial growth factor receptor (VEGF) tyrosine kinases involved in clinical trials concerning non-small cell lung cancer (NSCLC), breast, liver, and colorectal cancer as well as leukemia (38–40). Contrary to the Dabrafenib’s example, Linifanib is one of the rare examples where good modeling results are achievable despite low standard deviation of the AUC distribution (Fig. 8b). The high correlation achieved by the OT RF model mainly comes from its ability to accurately predict lowered AUC for three outlying, sensitive cell lines. The most decisive predictive feature in this model is the expression of FLT3 gene, which exhibits high over expression in these cell lines, with much higher mean 11.53 expression than the mean 3.30 for all cell lines in the training set. The expression of FLT3 ranks lower (11th) among features of the genome-wide model.

Similarly to Linifanib, Quizartinib is also characterized by low variation in the treatment response (Fig. 8c), and is also an FLT3 inhibitor. Quizartinib is tested in clinical trials for acute myeloid leukemia (AML) (41). The best biologically-driven model (PG RF) uses features related to genes present in drug’s target pathway (218 features), and the most important feature is expression of FLT3. The accurate prediction done by PG RF model for the single outlying, responsive sample (Fig. 8c) probably arises from the over-expression of FLT3 in that cell line (11.20 value for that feature in this sample vs. the mean of 3.26 for all training samples). Although expression of FLT3 also appears as the fourth most important feature in the genome-wide model, it is unable to correctly predict AUC for the responsive cell line, since the relative impact of FLT3 is much smaller. Overall, these three examples well show that feature selection can facilitate derivation of interpretable insights.

## Discussion

This work, to our knowledge, is the first comprehensive analysis of feature selection strategies for drug sensitivity prediction. Previous systematic assessments (13, 14) compared different modeling techniques and data types describing the cell lines, but did not comprehensively evaluate feature selection approaches. Similarly, although numerous modeling methods were developed specifically for the task of drug sensitivity prediction (7, 8), they were solely optimized for predictive power and not interpretability. If feature selection was applied at all, it was not driven by pre-existing biological knowledge, but performed using standard and often not robust selection techniques such as regularization (22).

Such comprehensive feature selection assessment is needed for several reasons. First of all, both feature selection driven by pre-existing biological knowledge and data driven selection have their advantages and disadvantages. Intuitively, selecting the features using *a priori* knowledge of the drug mode of action as a guideline should improve modeling. On the other hand, it is also restricting the available information for the model, and if the prior knowledge is wrong, may result in missing important dependencies. Given the vast number of features compared to the number of samples, the models with genome-wide data as features or ones with automated feature selection are badly ill-posed and prone to over-fitting. On the other hand, they are given the advantage of a larger number of samples (resulting in higher power) and access to more information, compared to the models with biologically driven features (Fig. 2). Second, as there is no obvious recipe for choosing the feature set for a particular drug, the in-depth comparative analysis of different feature selection strategies may suggest indications for the recommended type of features for drugs depending on their mode of action or knowledge of their target pathway. Finally, if the best performing feature set is small, each particular feature can be inspected and further evaluated as a potential biomarker for the drug. Here, different feature selection strategies driven by prior knowledge were compared to using genome-wide feature sets and the data-driven, automatic feature selection techniques across all analyzed drugs. We identified the best suited feature set for each drug and investigated them in the context of drugs’ target pathways. Finally, we evaluated the predictive power of different features types and inspected example drug-specific models in more detail. The entire assessment workflow aimed at the identification of such strategies that could deliver highly predictive, but also highly interpretable models, bringing insights about specific drugs that are informative for their application in precision medicine.

Our analysis provides extends insights into the drug response modeling aspects gained from the previous assessments. Jang *et al.* (14) concluded that the best modeling practice is to use elastic net or ridge regression with AUC as the response variable, with genome-wide gene expression features. Their assessment relied solely on correlation as model performance measure. Our analysis highlights AUC as a very difficult response variable, with the distribution strongly centered around value 1 for many of the compounds. With the actually interesting indicator of drug response being low AUC value, the model is given a difficult task to correctly learn this response from often very few examples. Interestingly, for these difficult cases, dummy models, predicting only the mean of the AUC for a given drug, can obtain surprisingly high correlation with the response. Thus, the actual performance evaluation measure should involve, in addition to correlation, also a comparison of the model RMSE to the dummy RMSE. Here, we implemented this comparison using Rel-RMSE, which facilitated filtering out degenerate models that do not perform better than the dummy model.

Both Jang *et al.* (14) and the DREAM challenge (13) assessments indicated that adding the features representing mutation and copy number status on top of genome-wide expression features did not improve the overall performance of modeling drug sensitivity (13, 14). This is likely due to the fact that gene expression is sometimes already reflecting genomic changes or tissue type. In contrast, our analysis shows that additional features corresponding to mutations are often significant predictors when they are evaluated as part of smaller feature set and are not vastly outnumbered by the gene expression features (for example, in the cases of Dabrafenib, PLX-4720, Nutlin-3a, SB590885 and Pelitinib).

## Conclusions

First, our results bring conclusions about feature selection strategies for drug sensitivity prediction. In general, the baseline genome-wide set of features or data-driven feature selection yields higher median predictive performance than biologically driven features. There are, however, multiple individual drugs, for which the feature selection driven by biological knowledge gives the best results, including models for the drugs with the top two performance scores. Moreover, feature selection driven by prior knowledge drastically reduces the number of features. At the same time, if the drop of performance in comparison with genome-wide models occurs, it is often only slight.

Second, the presented analysis gives important insights on how cancer cell lines react to different drugs. Drugs that are generally toxic or target general cellular mechanisms such as DNA replication or metabolism affect a relatively large proportion of cancer cell lines and thus have a wide response distribution. These compounds tend to be better modeled using genome-wide features, indicating that their effect on the cancer cells depends on a large spectrum of different cellular features. Conversely, for drugs targeting specific pathways, sensitivity distribution tends to be narrow, with most cells not responding at all and only a few interesting outliers of sensitive cells. For these compounds, high-level drug properties such as direct targets or target pathways allow to build highly predictive models with small numbers of interpretable features, such as Dabrafenib, Linifanib or Quizartinib. In particular, highly predictive models with an extremely low number of input features can be obtained, as in the cases of Linifanib, Afatinib, and GNF-2. Overall, this analysis shows the importance of using adequate feature selection strategies for each individual drug.

