## Supplementary Material for "Feature selection strategies for drug sensitivity prediction"

KRZYSZTOF KORAS, DILAFRUZ JURAEVA, JULIAN KREIS, JOHANNA MAZUR, EIKE STAUB, EWA SZCZUREK

### 1. Supplementary Methods

1.1. **Elastic net regression.** Elastic net regression belongs to the family of regularized linear regression models where the target value is expected to be a noisy linear combination of input features. The model introduces the regularization through adding a combination of $\ell_1$ and $\ell_2$ norms of it's coefficients to the loss function. Therefore, elastic net's optimization problem can be represented as:

$$\min_w \frac{1}{n}\|Xw - y\|_2^2 + \alpha\rho\|w\|_1 + \alpha(1-\rho)\|w\|_2^2$$

where $n$ is the number of samples, $X$ and $w$ represent the input data and coefficients vector, respectively. The amount and type of regularization are controlled by hyperparameters $\alpha$ and $\rho$, corresponding to *alpha* and *l1_ratio* arguments in scikit-learn implementation (1). These two parameters where tuned during cross-validation.

1.2. **Random forest regression.** Random forest is an ensemble method, which works by combining the outputs of several decision trees in order to make final predictions. In contrast to linear regression, decision trees is a non-parametric method which learns simple decision rules inferred from the data features. It's goal is to partition the input feature space such that samples with similar labels are grouped together. At each node $m$, corresponding data $Q$ is split into two subsets:

$$\begin{aligned} Q_{left}(\theta) &= (x, y)|x_j < t_m \\ Q_{right}(\theta) &= Q \setminus Q_{left} \end{aligned}$$

where $\theta$ is a candidate split consisting of a feature $j$ and corresponding threshold $t_m$ and $(x, y)$ represents training samples. Decision trees select parameters $j$ and $t_m$ which minimize the impurity of resulting subsets. The choice of a specific impurity function depends on application. In our analysis, we used mean squared error, which is common for regression tasks.

Decision trees have many advantages, but are also prone to create over-complex graphs which tend to overfit the data. In random forest, each tree is built from the bootstrap sample from the training set. Furthermore, the best split at each node is picked from a random subset of features. Such randomness, combined with averaging the predictions of single trees, helps to decrease the variance of the overall model. The hyperparameters we tuned when using random forests included (following scikit-learn notation): *n_estimators* – number of trees in the ensemble, *max_features* – maximum number of features considered when splitting the data, *max_depth* – maximum depth of the trees, *min_samples_split*

– minimum number of samples required to perform the split and *min_samples_leaf* – minimum number of samples allowed in a leaf node.

1.3. **Stability selection with lasso regression.** Stability selection (2) works by generating $N$ bootstrap samples of available data and using an underlying feature selection algorithm (in this case lasso regression) to determine which features are relevant for a given sample. For every generated sample, it fits the selection algorithm with a specified value of regularization parameter $\lambda$, which produces a selection set $\hat{S}_i^\lambda$ indicating which features to choose. Given selection sets from each sample, the empirical probability of choosing a particular feature $k$ can be computed as:

$$\hat{\Pi}_k^\lambda = \Pr[k \in \hat{S}^\lambda] = \frac{1}{N}\sum_{i=1}^{N} \mathbb{I}_{\{k \in \hat{S}_i^\lambda\}}$$

i.e. counting the number of times $k$ occurred as an important component for in the samples. This process is then repeated for several values of $\lambda$. The final stable set of relevant features can be then constructed as follows:

$$\hat{S}^{\text{stable}} = \{k : \max_{\lambda \in \Lambda} \hat{\Pi}_k^\lambda \geq \pi_{\text{thr}}\}$$

where $\Lambda$ is a set of all $\lambda$ values and $\pi_{\text{thr}}$ is a predefined probability threshold. In our work, we used scikit-learn compatible implementation of stability selection (3) combined with lasso regression, fitting for $N = 100$ samples with five different values of $\lambda$: $10^{-5}, 10^{-4}, 5 \cdot 10^{-4}, 10^{-3}$ and $10^{-2}$.

When applying automated stability selection, we first fitted the model using five different values of $\lambda$ and 100 bootstrap samples, which resulted in stability scores corresponding to every feature. We then iterated over predefined range (0, 1) of stability thresholds $\pi_{\text{thr}}$ with 0.025 increment, performing the whole modeling process with a corresponding number of features at each iteration using elastic net regression. This procedure was repeated for five random data splits. In order to establish the single best stability threshold for every compound, we averaged the results over data splits. The performance metrics used to evaluate the model were then the averages of metrics achieved with the chosen best threshold for every data split.

1.4. **Feature importance derived from random forest.** In a single decision tree, the depth of a feature used as a decision node represents the relative importance of that feature when predicting the target variable. Features present at the top of a tree contribute to the final prediction result for a bigger fraction of samples. The importance of a particular feature is also associated with the decrease of impurity when splitting the data using that feature (i.e. the bigger the importance, the bigger decrease in impurity measure). Therefore, the corresponding impurity decrease can be used to estimate the feature importance in a single tree (4). In random forests, this predictive ability of a given feature can be averaged over several trees to define a new metric, *Mean Decrease Impurity* (MDI) (5), which provides the feature importance estimate with reduced variance.

When using random forest for feature selection, after data extraction and hyperparameter tuning steps, we first trained the algorithm on the whole training set and extracted a vector with values representing the importance of all features. We then ranged over a grid of values $k$, each time performing the whole modeling process using random forests regression with $k$ most important features and recording the corresponding results. Similarly, as in the stability selection setting, one best value of $k$ was chosen by averaging the results over five data splits, and the corresponding performance metrics for best found $k$ were averaged in order to evaluate the model.

1.5. **Relative root mean squared error.** In each experimental setting, baseline, dummy model was deployed which always predicted the mean value of AUC in the training data. *Relative Root Mean Squared Error* (RelRMSE) is then defined as the fraction of the dummy model's RMSE on the test data and actual model's RMSE on the test data:

$$\text{RelRMSE} = \frac{\text{RMSE}_{dummy}}{\text{RMSE}_{model}}$$

i.e. better performance corresponds to bigger RelRMSE metric, with a baseline score of 1.
