## Supplementary Table S1 for "Feature selection strategies for drug sensitivity prediction"

Gene expression signatures along with corresponding number of genes and reference. For signatures with missing references, list of genes is provided below the table.

| Signature Name | Genes | PMID |
|---|---|---|
| Kannengiesser.BRAF.signature | 25 | 19383316 |
| IFN_signature | 32 | |
| KinetochoreNet | 33 | |
| B_cell_signature_IRIS | 90 | |
| T_cell_signature_IRIS | 14 | |
| DNAsynthesisFuncNet | 64 | |
| Chi.Hypoxia.BrownVanDeVijver | 18 | 16417408 |
| Breast_ERBB2subtype.Calza2006 | 8 | 16846532 |
| Breast_BASALsubtype.Calza2006 | 17 | 16846532 |
| Breast_LuminalBsubtype.Calza2006 | 9 | 16846532 |
| ER_pos_BreastCa.Abba2005 | 9 | 15762987 |
| TGFbeta.EarlyUp.Verrecchia2001 | 50 | 11279127 |
| TNFa_NFkB_pw_response.Tian2005 | 20 | 15722553 |
| B_cell_signature.Newell2010.markers | 18 | 20501946 |
| medulloblastoma.IFN.up.Staub2012 | 10 | 22937184 |
| medulloblastoma.PRF.up.Staub2012 | 21 | 22937185 |
| BreastCaCellCycleSig.Dai2005 | 36 | 15899795 |
| BreastStromaMetagene.Farmer2005 | 50 | 15897907 |
| AKT.up.Creighton2007 | 57 | 17213801 |
| EMT.BreastCa.Lien2007 | 33 | 17603561 |
| ClassicalPancreasCa.Collison2011 | 22 | 21460849 |
| Phillips2006.ProNeural | 15 | 16530701 |
| Phillips2006.Prolif | 5 | 16530702 |
| Phillips2006.Mesenchymal | 15 | 16530703 |
| Lottaz2010.TypeII.CSC | 21 | 20145155 |
| Lottaz2010.TypeI.CSC | 8 | 20145156 |
| Freije2004.GBM.prognGroup.HC2A | 10 | 15374963 |
| Freije2004.GBM.prognGroup.HC2B | 15 | 15374964 |
| CDK8_genomic_neighbors | 6 | |
| Liang2005.GBM-Hypoxia | 21 | 15827124 |
| Liang2005.GBM-ECM | 19 | 15827125 |
| Liang2005.GBM-Prolif | 22 | 15827126 |
| Farmer2005.BreastCa.Clu1.IFN_Tcell_Bcell | 44 | 15897900 |
| Farmer2005.BreastCa.Clu2.prolif_8qamplicon | 33 | 15897901 |
| Farmer2005.BreastCa.Clu3.apocrine_basal_hypoxia | 15 | 15897902 |
| Farmer2005.BreastCa.Clu5.17q21_32amplicon | 17 | 15897903 |
| Farmer2005.BreastCa.Clu6.luminal | 16 | 15897904 |
| Farmer2005.BreastCa.Clu7.apocrine_luminal | 20 | 15897905 |
| Farmer2005.BreastCa.Clu8.ERBB2amplicon | 7 | 15897906 |
| Farmer2005.BreastCa.Clu4.stroma | 19 | 15897907 |
| EMTup.QiagenPCRarray | 36 | |
| Kuner2009.LungSCCvsAD | 10 | 18486272 |

| | | |
|---|---|---|
| Cytosolic.Ribosomal.Proteins | 68 | |
| Sadanandam2013_InflammatoryCRC | 16 | 23584089 |
| Sadanandam2013_StemLikeCRC | 164 | 23584093 |
| Wilkerson2010_LungSqCC_PrimitiveSubtype | 30 | 20643781 |
| Wilkerson2010_LungSqCC_ClassicalSubtype | 18 | 20643781 |
| Wilkerson2010_LungSqCC_SecretorySubtype | 24 | 20643781 |
| Wilkerson2010_LungSqCC_BasalSubtype | 17 | 20643781 |
| SomaLogic_PlateletActivationInPlasma | 15 | |
| ECM_qPCR_panel_Qiagen | 83 | |
| Feng2006_IFN_5gSig | 5 | 16947629 |
| Huang2012_EMTdown | 265 | 23165231 |
| Huang2012_EMTup | 229 | 23165231 |
| Huang2012_MED12kd_MEKiResist_54genes | 54 | 23165231 |
| VanDerFlier2007_WNTsig_DwAftDomNegTCFexpr | 15 | 17320548 |
| DeSousa2013_CCS3_Serrated_46genes | 46 | 23584092 |
| colon.MEKiSS.s32_down | 25 | |
| colon.MEKiSSsecreted.s32_down | 25 | |
| KRAS.dependency.signature_up | 33 | |
| Dry.signature_up | 17 | |
| Dry.signature_down | 13 | |
| EMT_LiteratureMarkers_up | 5 | |
| EMT.Taube.Weinberg.GSE24202_up | 93 | 20713713 |
| EMT.Taube.Weinberg.GSE24202_down | 156 | 20713713 |
| HippoPWup.YAPtransfection.GSE10196_down | 43 | 18413746 |
| BreastCa.PoorPrognosisConsensus.Teschendorff2006_up | 15 | 17076897 |
| p53-mut.Miller2005_up | 5 | 16141321 |
| PTENlossBreastCa.Saal2007_up | 112 | 18066063 |
| p53mut.BreastCa.Troester2006_up | 32 | 17150101 |
| Epithelial.markers.Literature_up | 34 | |
| Verhaak2010.Proneural_up | 137 | 20129251 |
| Verhaak2010.Proneural_down | 73 | 20129251 |
| Verhaak2010.Mesenchymal_down | 46 | 20129251 |
| Prat_ClaudinLow_up | 425 | 20813035 |
| Prat_ClaudinLow_down | 357 | 20813035 |
| S7_mDCs_plus_Monocytes | 15 | |
| S10_B_cells | 50 | |
| S12_T_cells_aCD3_aCD28_activated | 50 | |
| Bennett2003.SLE.granulopoiesis.signature | 20 | 12642603 |
| Bennett2003.SLE.ifn.sig | 26 | 12642603 |
| Chaussabel2008_M1.2_ifn.sig | 27 | 18631455 |
| Chaussabel2008_M3.4_ifn.sig | 53 | 18631455 |
| Chaussabel2008_M5.12_ifn.sig | 59 | 18631455 |
| Rice2013_IFNsig_6g | 6 | 24183309 |
| Walsh2007_IFNsig_6g | 6 | 17968926 |
| PlasmaCell_signature_24g | 24 | |
| Budinska2013.deregulated.in.CRC | 27 | 23836465 |
| Budinska2013.chromosome20q.CRC | 33 | 23836465 |

| | | |
|---|---|---|
| Budinska2013.proliferation | 83 | 23836465 |
| Budinska2013.colon.crypt.markers.CRC | 16 | 23836465 |
| Budinska2013.EMT.stroma.CRC | 310 | 23836465 |
| Budinska2013.immune.response.CRC | 102 | 23836465 |
| Wright2003_ABC_DiffuseLargeBcellCarcinoma | 21 | 12900505 |
| Speers2015_BrCa_radioresistance_down | 28 | 25904749 |
| Lehmann.2011.BL2.up.refined | 23 | 21633166 |
| Lehmann.2011.IM.up.refined | 174 | 21633166 |
| IFNsig.Staub2015 | 7 | |
| Walter2013_HNCa_atypical_high | 132 | 23451093 |
| Walter2013_HNCa_classical_high | 62 | 23451093 |
| Walter2013_HNCa_mesenchymal_high | 246 | 23451093 |
| MasqueSoler2013_DLBCL_nonMolBL | 4 | 24030260 |
| MasqueSoler2013_DLBCL_molBL | 6 | 24030260 |
| MasqueSoler2013_DLBCL_ABC | 11 | 24030260 |
| Scott2014_DLBCL_ABC | 8 | 24398326 |
| YAP_target_genes | 8 | |
| DNArepairScore_high_Kang2012 | 19 | 22505474 |
| DNArepairScore_low_Kang2012 | 4 | 22505474 |
| DDR_genes | 43 | |
| Mulligan2014_DDRD_up | 24 | 24402422 |
| Mulligan2014_DDRD_Group3 | 7 | 24402422 |
| Mulligan2014_DDRD_Group4 | 10 | 24402422 |
| Hou2010_LuCa_SCC | 47 | 20421987 |
| Hou2010_LuCa_ADC | 5 | 20421987 |
| radio_sensitivity_genes_up_BMC_Kim2012 | 10 | 22846430 |
| radio_sensitivity_genes_down_BMC_Kim2012 | 21 | 22846430 |
| radioresistance_PNAS_Khodarev2004 | 51 | 14755057 |
| YAP_20gSig_Staub2016 | 20 | |
| YAP_23gSig_Staub2016 | 23 | |
| YAP_48gSig_Staub2016 | 48 | |
| DDR_Alt-NHEJ | 4 | |
| DDR_FA (Fanconi anemia pathway) | 37 | |
| DDR_HR (Homologous Recombination) | 52 | |
| DDR_MMR | 26 | |
| Platinum_sensitivity_JNCI2012 | 23 | 22505474 |
| Topotecan_sig_Pitroda_2014 | 12 | 24670686 |
| RPS_Pitroda_2014 | 4 | 28341751 |
| PARPi_Deamon_2012 | 7 | 22875744 |

**IFN_signature**

ADAR, CCR2, CIC, CXCL10, FADS1, FCGR1A, IFI27, IFI44, IFI6, IFIT1, IFIT2, IFIT3, IFIT5, IL1RN, ISG15, LGALS3BP, LY6E, MARCKS, MX1, MX2, OAS1, OAS2, OAS3, OASL, PLSCR1, RASGEF1B, RNASE2, SERPING1, SOCKS1, STAT1, TNFSF10, XAF1

**KinetochoreNet**

BUB1, BUB1B, CALCOCO1, KNL1, CBX3, CBX5, CENPE, CENPH, CETN3, DSN1, E2F1, E2F4, FOXO1, HNF1A, HNF4A, KDM5B, KLHL12, MIS12, NDC80, NEK2, NSL1, NUF2, PMF1, PSMC2, RB1, SMC1A, SPC24, SPC25, UBR5, USHBP1, ZW10, ZWINT

**B_cell_signature_IRIS**
ALG5, AMPD1, AP1, B4GALT3, TNFRSF13C, BANK1, BCL11A, TNFRSF17, FAM129C, BLK, BLNK, BMP8B, STAP1, VCPKMT, MYDGF, EDEM2, C21orf83B, CD19, MS4A1, CD79A, CD79B, KLF6, CPNE5, CXCR5, DDOST, DKFZp667L0210, DTNB, EAF2, EIF2AK3, ELL2, ERN1, PDIA4, EST, FBH1, FCRL1, FCRL2, FKBP11, TENT5C, TXNDC15, TMEM156, EME1, DERL3, FCRLA, GNG7, GPRC5D, PLPP5, FCRL5, SPCS2, FAM30A, SEL1L3, KLHL14, LOC220213, LOC51061, LZTFL1, MAN1A1, MANEA, NXPE3, MT-ND6, NLRP7, NGLY1, OSBPL10, PACAP, PAX5, PC4, PNOC, POU2AF1, QRSL1, RALGPS2, RPN1, SCFD1, SEC24A, KDM5D, SPATS2, SPIB, SSR1, HSPA13, TCF3, TCL1A, SEC62, TLR10, HSP90B1, TRAM1, TRAM2, TXNDC5, UBE2G1, UBE2J1, Ufm1, EZR, VPREB3, WNT10A

**T_cell_signature_IRIS**
BCL11B, CD3D, CD3E, CD3G, CD5, CD6, CD8A, CD8B, CTLA4, CXCR6, IL17F, IL22, IL9, TRA

**DNAsynthesisFuncNet**
APEX1, CCNA2, CCND1, CCND2, CCND3, CCNE2, CCNG1, CDC25C, CDK2, CDK4, CDK5, CDK6, CDK7, CDKN1A, CDKN1B, CDKN1C, CDT1, CHAF1A, CHTF18, DNMT1, DUSP1, EP300, FEN1, FZR1, HDAC1, HELB, HNRNPA1, PCLAF, LIG1, MRE11, NBN, NEIL1, NEIL2, PARP1, PCNA, POLA1, POLB, POLD1, POLD2, POLD3, POLD4, POLDIP2, POLE, POLH, POLI, POLM, RAD18, RAD9A, RBL1, RBL2, RECQL4, REV1, RFC2, RFC3, RPA1, RUVBL2, SKP2, TERT, TYMS, WRN, XRCC1, XRCC5, XRCC6, YBX1

**CDK8_genomic_neighbors**
NUP58, RNF6, CDK8, GPR12, USP12, RASL11

**EMTup.QiagenPCRarray**
AHNAK, BMP1, CALD1, CAMK2N1, CDH2, COL1A2, COL3A1, COL5A2, FN1, FOXC2, GNG11, GSC, IGFBP4, ITGA5, ITGAV, MMP2, MMP3, MMP9, MSN, SERPINE1, SNAI1, SNAI2, SNAI3, SOX10, SPARC, STEAP1, TCF4, TIMP1, TMEFF1, TMEM132A, TWIST1, VCAN, VIM, VPS13A, WNT5A, WNT5B

**Cytosolic.Ribosomal.Proteins**
RPL10, RPL10A, RPL11, RPL12, RPL13, RPL13A, RPL13P5, RPL14, RPL15, RPL17, RPL18, RPL18A, RPL21, RPL22, RPL23, RPL24, RPL26, RPL27A, RPL28, RPL29, RPL3, RPL30, RPL31, RPL32, RPL35, RPL35A, RPL36, RPL37, RPL37A, RPL39L, RPL3L, RPL4, RPL5, RPL6, RPL7, RPL7A, RPL8, RPLP1, RPLP2, RPS10, RPS11, RPS12, RPS13, RPS14, RPS15, RPS15A, RPS16, RPS17, RPS18, RPS19, RPS2, RPS20, RPS21, RPS25, RPS26, RPS27, RPS27A, RPS27L, RPS28, RPS29, RPS3, RPS3A, RPS4Y1, RPS5, RPS6, RPS7, RPS8, RPS9

**SomaLogic_PlateletActivationInPlasma**
BDNF, TIMP3, CCL5, MMP9, PF4, ANGPT1, MDK, SERPINE1, SPARC, APP, CTSA, SERPINE2, DKK4, THBS1, PDGFB

**ECM_qPCR_panel_Qiagen**
ADAMTS13, ADAMTS8, MMP1, MMP10, MMP11, MMP12, MMP13, MMP14, MMP15, MMP16, MMP2, MMP3, MMP7, MMP8, MMP9, SPG7, TIMP1, CD44, CDH1, CLEC3B, CNTN1, COL11A1, COL12A1, COL14A1, COL15A1, COL16A1, COL1A1, COL4A2, COL5A1, COL6A1, COL6A2, COL7A1, COL8A1, FN1, ANOS1, THBS1, TIMP2, TIMP3, CTGF, CTNNA1, CTNNB1, CTNND1, CTNND2, VCAN, ECM1, HAS1, SPP1, TGFBI, THBS2, THBS3, TNC, VTN, ICAM1, ITGA1, ITGA2, ITGA3, ITGA4, ITGA5, ITGA6, ITGA7, ITGA8, ITGAL, ITGAM, ITGAV, ITGB1, ITGB2, ITGB3, ITGB4, ITGB5, LAMA1, LAMA2, LAMA3, LAMB1, LAMB3, LAMC1, NCAM1, PECAM1, SELE, SELL, SELP, SGCE, SPARC, VCAM1

**colon.MEKiSS.s32_down**
C3, MMP9, DCN, SERPINF1, MGP, C1S, CDH11, SNORD114-3, ISLR, CTSK, MYH11, IL3RA, SULF1, ANTXR1, LUM, FBLN5, THBS2, C1R, ACTA2, IGFBP5, MXRA5, APOD, GUCY1A1, BGN, CRISPLD2

**colon.MEKiSSsecreted.s32_down**

C3, MMP9, SERPINF1, DCN, IGFBP5, ISLR, MMP2, LUM, MGP, FBLN5, AEBP1, A2M, MFAP4, ASPN, SPARCL1, OLFML2B, CRISPLD2, TIMP3, BGN, SRPX2, COL6A1, APOD, CXCL12, SPARC, AOAH

**KRAS.dependency.signature_up**
SYK, ESRP1, ST14, TMEM30B, SPINT1, RAB25, KDF1, GRHL2, GALNT3, SCNN1A, MPZL2, ITGB6, IRF6, INPP4B, PCDH1, C6orf141, HS3ST1, CDS1, DNAJA4, F11R, PROM2, CLDN7, C1orf116, SCEL, SCIN, S100A14, ANKRD22, MAL2, EHF, MSRB3, INAVA, TTC9, DENND1C

**Dry.signature_up**
ZNF106, PROS1, LZTS1, TRIB2, DUSP4, ETV4, ETV5, DUSP6, PHLDA1, SPRY2, ELF1, LGALS3, FXYD5, S100A6, SERPINB1, SLCO4A1, MAP2K3

**Dry.signature_down**
IL6, CD274, G0S2, STAC, COL5A1, COL12A1, SERPINE1, CRIM1, LOX, GPR176, FZD2, BASP1, CLU

**EMT_LiteratureMarkers_up**
SNAI1, SNAI2, TWIST1, VIM, CDH2

**Epithelial.markers.Literature_up**
SH2D3A, JUP, RAB25, CDH1, LSR, SPINT2, DDR1, GRHL2, GALNT3, CDS1, MAL2, CRB3, EPCAM, LLGL2, LAD1, TMEM125, PRSS8, SFN, ELF3, C1orf116, OCLN, PPL, INAVA, MAP7, ARHGEF5, S100A14, CDH3, MACC1, CHMP4C, HOOK1, CBLC, DSC2, PLS1, MAP3K9

**S7_mDCs_plus_Monocytes**
GCH1, CMPK2, ISG15, CXCL10, CXCL9, IFI6, EPSTI1, IL15RA, BATF3, IL15, APOL3, IFI44, IFIH1, ZNRF1, LOC100506459

**S10_B_cells**
BLK, CD79A, CD79B, CXCR5, FCRL1, P2RX5, FAM30A, POU2AF1, VPREB3, PCDH9, FCRL5, QSOX2, FCRLA, KLHL14, SGCE, IGHM, DSP, CCDC191, PAX5, PEG10, IGLJ3, STAG3, BTLA, LOC100130458, EML6, SLC38A11, CD19, CPNE5, CD24, SNX22, CD22, STRBP, CD200, PIK3C2B, STAP1, SYBU, CNTNAP2, IGLL3P, LARGE2, LINC00926, HIP1R, DTX1, LOC100507616, PLEKHG1, MACROD2, ABCB4, GGA2, IGHD, HLA-DOB, PLPP5

**S12_T_cells_aCD3_aCD28_activated**
CCNA2, PBK, PTTG1, SAAL1, ZWINT, UBE2T, MAD2L1, CXCR6, UTP15, DEPDC1B, MELK, CDCA7, NCAPG2, CENPH, FIGNL1, NDC80, CDKN3, ZC3HAV1L, POLE2, LAG3, SLC25A17, CENPM, COL6A3, HPGD, TIPIN, RMI2, NUF2, NCAPH, RTTN, RRM2, MCM2, PCLAF, TMEM200A, WDR89, TMEM135, NPM3, ZBTB9, FAM83D, TYMS, CENPK, GPR171, CHAF1B, NECTIN3, GRPEL2, KLC2, DCLRE1A, SLC9B2, C5orf30, WEE1, ABCD2

**PlasmaCell_signature_24g**
PDIA6, PRDM1, MAN1A2, RABAC1, CAV1, IGF1, HYOU1, HSPA13, CD38, ELL2, UAP1, SDC1, B9D1, STT3A, IGLV1.44, MYDGF, WFS1, PDIA4, RRBP1, GFPT1, TNFRSF17, MAN1A1, HERPUD1, RWDD2A

**IFNsig.Staub2015**
IFIT3, IFIT2, IFIT1, IFI44, IFI44L, OASL, OAS3

**YAP_target_genes**
CTGF, CYR61, ANKRD1, FOSL1, ACTN1, PDLIM7, AXL, ODC1

**DDR_genes**
AKT1, ATM, ATR, BAP1, BARD1, BRCA1, BRCA2, BRIP1, CDK12, CHEK1, CHEK2, CTNNB1, ERCC4, ABRAXAS1, FANCA, FANCD2, FANCE, FANCI, FANCL, KRAS, MLH1, MRE11, MSH2, MSH6, MUTYH, NBN, PALB2, PIK3CA, PPP2R2A, PTEN, RAD50, RAD51, RAD51B, RAD51C, RAD51D, RAD52, RAD54B, RAD54L, RPA1, TP53, TP53BP1, XRCC2, XRCC3

**YAP_20gSig_Staub2016**

AMOTL2, GPRC5A, TLCD2, TJP1, BCL9L, AJUBA, SDC4, RBMS2, CRIM1, TNFRSF12A, EPHA2, FHL2, FOSL1, ANXA2, CYR61, MYOF, CAVIN1, RND3, LOC100288911, NTN4

**YAP_23gSig_Staub2016**
AJUBA, AMOTL2, ANXA2, AXL, BCL9L, BOK, CRIM1, CYR61, DCBLD2, EPHA2, FHL2, FOSL1, GPRC5A, MYOF, NTN4, CAVIN1, RBMS2, RND3, SDC4, TJP1, TLCD2, TNFRSF12A, YAP1

**YAP_48gSig_Staub2016**
AMOTL2, GPRC5A, TLCD2, TJP1, BCL9L, AJUBA, SDC4, RBMS2, CRIM1, TNFRSF12A, EPHA2, FHL2, FOSL1, ANXA2, CYR61, MYOF, CAVIN1, RND3, LOC100288911, NTN4, ITGA3, DCBLD2, AXL, CAV1, AHNAK2, TGFBI, MT2A, CAV2, TNFAIP1, RTN4, TIMP2, YAP1, ERBB2, TUFT1, EDN1, CLIC3, ATP8B1, SSH3, C6orf132, DSP, KRT19, SERINC2, KIAA1522, BOK, RHOD, PPP1R13L, F3, RHPN2

**DDR_Alt-NHEJ**
LIG1, LIG3, PARP1, XRCC1

**DDR_FA (Fanconi anemia pathway)**
CENPS, BARD1, BLM, BRCA1, BRCA2, BABAM2, BRIP1, ABRAXAS1, DNA2, FAAP100, FAAP24, FAN1, FANCA, FANCB, FANCC, FANCD2, FANCE, FANCF, FANCG, FANCI, FANCL, FANCM, HELQ, HES1, KAT5, PALB2, RAD51, RAD51C, RMI2, STRA13, BHLHE40, TELO2, TOP3A, TOP3B, UBE2T, USP1, WDR48

**DDR_HR (Homologous Recombination)**
BLM, BRCA1, BRCA2, EID3, EME1, EME2, GEN1, H2AFX, HELQ, HFM1, KAT5, MRE11, MUS81, NBN, NSMCE3, NFATC2IP, NSMCE1, NSMCE2, NSMCE4A, PARG, PAXIP1, PPP4C, PPP4R1, PPP4R2, PPP4R4, RAD50, RAD51, RAD51B, RAD51C, RAD51D, RAD52, RAD54B, RAD54L, RDM1, RECQL, RECQL4, RECQL5, RMI2, RPA1, RPA2, RPA3, RPA4, SEM1, SLX1A, SLX4, SMC5, SMC6, SPO11, TOP3A, TOP3B, UIMC1, WRN

**DDR_MMR**
EXO1, HMGB1, LIG1, MLH1, MLH3, MSH2, MSH3, MSH4, MSH5, MSH6, PCNA, PMS1, PMS2, POLD1, POLD2, POLD3, POLD4, RFC1, RFC2, RFC3, RFC4, RFC5, RPA1, RPA2, RPA3, RPA4
